# A bite force database of 654 insect species

**DOI:** 10.1101/2022.01.21.477193

**Authors:** Peter Thomas Rühr, Carina Edel, Melina Frenzel, Alexander Blanke

## Abstract

Bite force is a decisive performance trait in animals because it plays a role for numerous life history components such as food consumption, inter- and intraspecific interactions, and reproductive success. Bite force has been studied across a wide range of vertebrate species, but only for 20 species of insects, the most speciose animal lineage. Here we present the insect bite force database with bite force measurements for 654 insect species covering 111 families and 13 orders with body lengths ranging from 4.2 - 180.1 mm. In total we recorded 1906 bite force series from 1290 specimens, and, in addition, present basal head, body, and wing metrics. As such, the database will facilitate a wide range of studies on the characteristics, predictors, and macroevolution of bite force in the largest clade of the animal kingdom and may serve as a basis to further our understanding of macroevolutionary processes in relation to bite force across all biting metazoans.

## Background & Summary

Bite force is a performance trait which may decide on an animal’s ability to acquire food, win inter- and intra-specific fights and successfully reproduce^1–5^. In vertebrates, maximum bite forces are well studied across a wide diversity of taxa such as bony fishes^6,7^, crocodilians (e.g. ^8^), birds (e.g. ^9^), turtles (e.g. ^10^), squamates (e.g. ^11,12^), frogs^13^, marsupials^14^, and mammals (e.g. ^15–18^).

Fundamental knowledge on the variation, predictors and evolution of bite forces within the omnipresent insects is, however, lacking. Even though more than half a million insect species belong to orders that possess biting-chewing mouthparts^19,20^, existing literature only yields bite force measurements on five dragonflies^21,22^, one cockroach^23^, and 14 beetles^24,25^. This is despite the fact that biting-chewing insects include the most destructive plant eating animals and occupy crucial roles in the world’s ecosystems as soil-building detritivores^26^.

So far, measuring bite forces of insects was hampered by their small size, but the recently published measurement setup “forceX” ^27^ overcame this limitation to some extent by allowing minimally invasive *in vivo* bite force measurements of animals with gape sizes more than ten times smaller than previous setups (e.g. ^28^). Using forceX, we measured bite forces of 654 insect species from 111 families and 13 orders, collected on four continents and from numerous breeding cultures. Instead of gathering maximum force values only, as most previous bite force studies have done (but see ^25,23,21,22,29^), we also recorded force curves. In addition, the bite force database contains head, wing and body metrics of each specimen to assess morphological predictors for bite force in insects. Thus, the database will facilitate investigations on the macroevolution of maximum bite force, bite lengths, bite frequencies, muscle activation patterns, and bite curve shapes across the megadiverse insects and will facilitate comparisons with all biting metazoan taxa.

## Methods

### Collection and material

A total of 1290 insect specimens representing 654 species from 111 families and 13 orders were collected in Australia, China, Denmark, France, Germany, Greece, Panama and Slovenia using light traps, insect nets, pitfall traps, or directly by hand. All specimens were collected under the respective regulations in effect (see Acknowledgements). Additionally, we measured specimens from numerous scientific, private and commercial insect breeders and traders (Online-only Table 1 and Acknowledgements).

### Size measurements

Head width, head length, head height, thorax width, forewing length and body length measurements were performed to the nearest 0.01 mm using a digital caliper (77001, Wentronic GmbH, Braunschweig, Germany). For the head width, the longest distance from left to right was measured, including protruding eyes if applicable (Fig. 1a). Head height in orthognathous insects was measured from the clypeo-labral ridge to the dorsal end of the head (Fig. 1a,b). In prognathous insects, head length was measured from the clypeo-labral ridge to the posterior end of the head (Fig. 1c). Thorax width was measured on the prothorax (Fig. 1d) and excluded lateral protrusions as found e.g. in many cockroaches and praying mantises. Body length measurements excluded cerci, ovipositors, or other abdominal appendages (Fig. 1e).

**Fig. 1:**
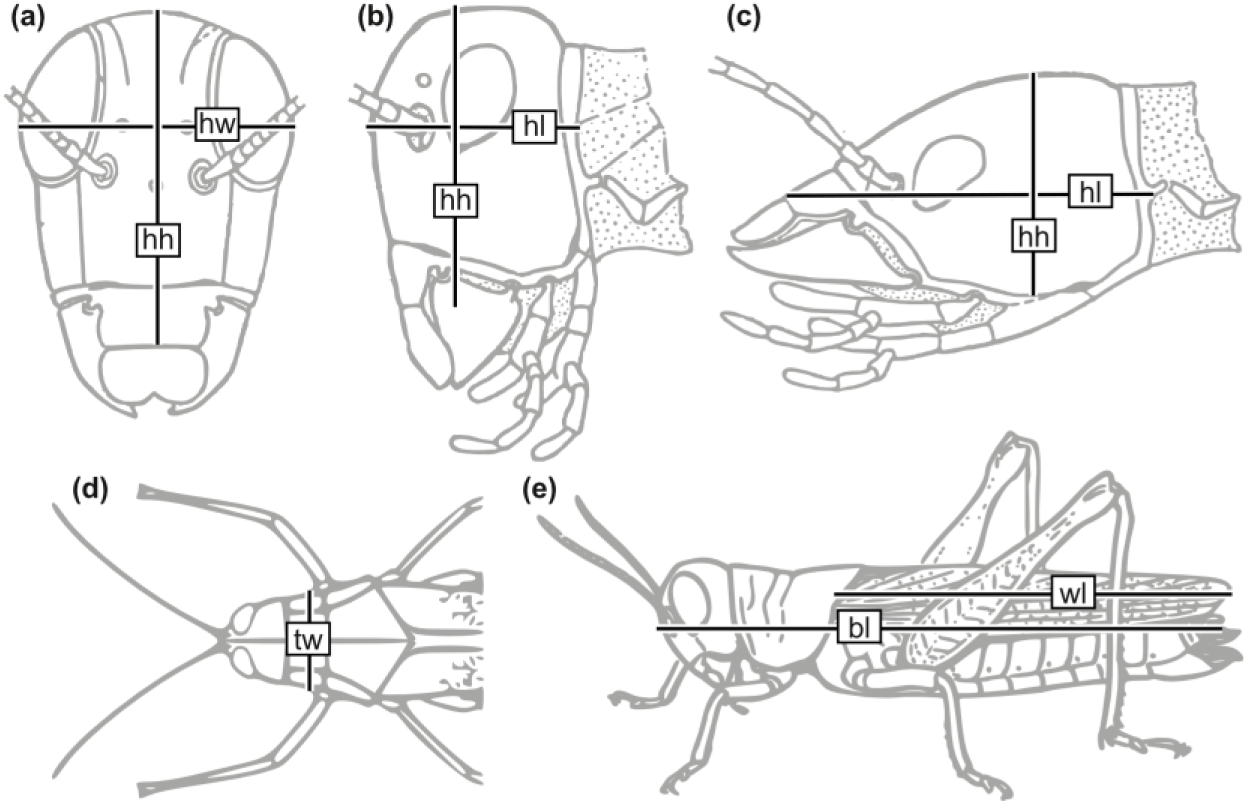
Insect length measurements. **(a-b)** Head of an orthognathous insect in frontal (a) and lateral view (b). **(c)** Head of prognathous insect in lateral view. **(d)** Frontal part of an orthognathous insect in dorsal view. **(e)** Orthognathous insect habitus in lateral view. Abbreviations: **bl**, body length; **hh**, head height; **hl**, head length; **hw**, head width; **wt**, thorax width; **wl**, forewing length. a,b,c after Snodgrass^42^; e after Snodgrass^43^.

### Bite force measurements

All measurements were carried out with the metal-turned version of the forceX setup as described in ^27^. In short, life and conscious animals were held between two fingers, rotated by 90° along their body axis and allowed to voluntarily bite on the tip elements of the forceX. Different tip element designs^27^ and distances between them were used to accommodate different animal gape sizes. During measurements, animals were observed through the Junior Stereo 3D microscope (Bresser GmbH, Rhede, Germany) that is part of the forceX setup to ensure that gape sizes are suitable and that the insects bite at the edge of the tip elements so that the ratio of the forceX lever remains at a constant 0.538^27,30^. We also checked if the animals bit with the distal-most incisivi of their mandibles to ensure that measurements remain comparable^30^. Non-distal bites or wrongly placed bites on the tip elements were discarded. If animals did not start biting by themselves, we used the tip element protrusions to insert the tip elements between the mandibles and/or used a fine brush to touch the animal’s cerci, head or abdomen^23^. Amplified analogue voltage signals were converted to a digital signal by a 12-bit USB data acquisition device (U3-HV, LabJack Corporation, Lakewood, Colorado, US) and recorded with the LJStreamUD v1.19 measurement software (LabJack Corporation) on a computer.

### Data curation

Subsequent data curation was performed in the software environment ‘R’ v.4.03^31^ using the package ‘forceR’ v.1.0.0^27^. Since the forceR package was written to analyse data generated with the forceX setup, we used, if not stated otherwise, the default settings of the package functions. First, time series were converted from the output format of LJStreamUD to a *.csv file containing only a time and a voltage column (without changing measurement values) using the forceR function ‘convert_measurement()’. Then, all measurements were manually cropped using ‘crop_measurement()’ to exclude regions without bite data at the beginning and end of each measurement. Next, ‘amp_drift_corr()’ was used to correct for the logarithmic drift of the analogue charge amplifier (see Rühr and Blanke ^27^ for details). When using the high amplification setting (20 V/N) to amplify the miniscule voltage signals of the piezoelectric force transducer at small bite forces, the zero-voltage-line (‘baseline’) may drift notably during a measurement. Therefore, a PDF file depicting all input raw data and their amplifier drift-corrected data graphs (available at Zenodo, s. Data Records) was visually inspected, and, if necessary, the function ‘baseline_corr()’ was used in its automatic mode to correct for this drift. In some of these cases, however, especially when the test animals showed long, plateau-like bite curve shapes, the automatic mode of ‘baseline_corr()’ failed to find the baseline, and the manual mode was used. All corrections can be retraced in the PDF file and reproduced using the log files that were created during corrections and which are stored at Zenodo. With the function ‘reduce_frq()’ we then reduced the sampling rate of all time series to 200 Hz, a value found to be sufficient to represent insect bite force curves^25,23,22,22^ to reduce the amount of data for further analyses. As a last curating step, voltage values were converted into force data [N] with the forceR function ‘y_to_force()’ that considers the amplification level of each measurement and the lever mechanics of the measurement system. Online-only Table 1 shows all measurement settings, taxonomic classifications and information on which correction procedures have been performed on which measurements.

### Maximum force value extraction for specimens and species

To extract maximum force values of each specimen and each species and calculate the standard deviations of these values we used the function ‘summarize_measurements()’ of forceR and custom code, heavily on the packages ‘dplyr’ v.1.0.7^32^. We then plotted the log10-transformed average maximum bite force per specimen (grey dots in Fig. 2) and per species (black dots) against the log10-transformed average body length (Fig. 2a) and head width (Fig. 2b) using ‘ggplot2’ v.3.3.5^33^ and ‘ggExtra’ v.0.9^34^. Linear regressions through the log10-transformed species-wise data showed significant allometric relationships between body size and head width (*p* < 0.001) with explanatory values of *r^2^* = 0.43 and *r^2^* = 0.56, respectively. Due to the expected logarithmic releationship between size and bite force^35^, means were calculated as geometric means. Calculations with the regular mean, however, yielded similar results (*p* 0.001, *r^2^* = 0.44 and *r^2^* = 0.56; Supplementary Figure 1).

**Fig. 2:**
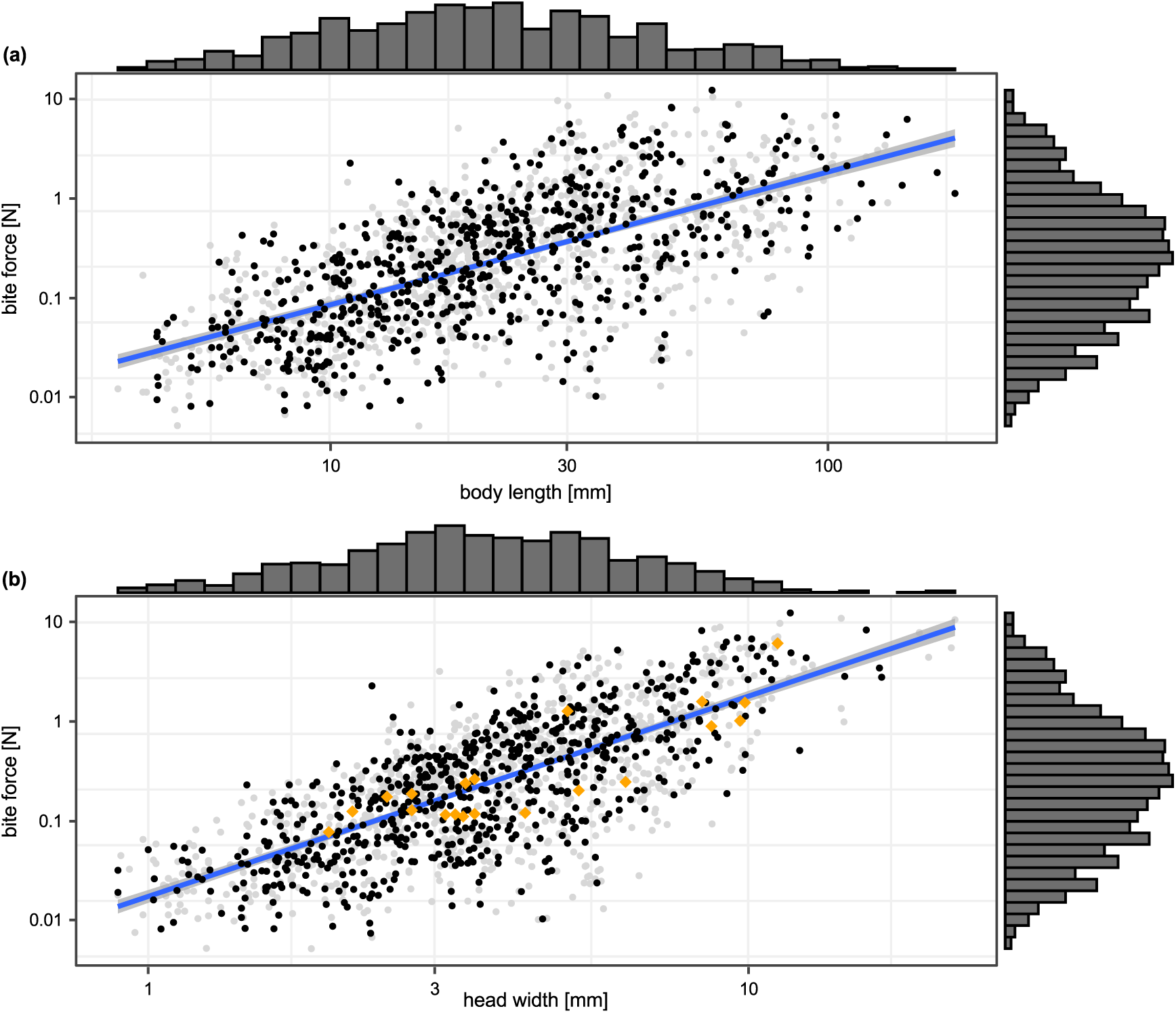
Maximum bite force against body length (a) and head width (b). Grey dots show geometric means of all maximum bite forces per specimen, black dots show geometric means of all length measurements and maximum bite forces of all specimens per species. Marginal histograms at the x- and y-axes show mean size and mean bite force distribution per specimen, respectively. Regression lines and coefficients refer to log10-linear models of species-wise bite force against body length (a) or head width (b). Orange diamonds in (b) show bite force measurements available in previous literature. All axes are log10-transformed.

### Comparison to previous insect bite force measurements

Previous studies on insect bite forces covered maximum bite force values for 20 species^24,23,21,22^. To check if these measurements follow similar allometric slopes as our data, we extracted all available insect bite force data from the literature and added them to the scatterplot in Figure 2b. We then tested if our data and the literature data differ in their allometric slopes by comparing a linear model with the null hypothesis of different slopes (log10(bite.force)~ log10(head.width) * source) versus a linear model with the null hypothesis of common slopes (log10(bite.force) ~ log10(head.width) + source). Both model fits were compared with an ANOVA to find out if they differ significantly.

### Assessment of geographical coverage

Climate zone data (Köppen–Geiger classification system^36–38^) was gathered for each species based on the GPS coordinates of its collection localities (Online-only Table 1) with the function ‘LookupCZ()’ of the R package ‘kgc’ v.1.0.0.2^39^. Percentages of species in the database for each country and climate zone were calculated.

### Assessment of phylogenetic coverage

To assess the phylogenetic coverage of the bite force database we compared the number of species with database entries to the number of species listed by the Open Tree of Life^40^, accessed on 2022/02/05 with the function ‘tol_node_info()’ of the package ‘rotl’ v.3.0.11^41^. Comparisons were carried out for all insect orders and families that are present in the bite force database.

## Supporting information

Online-only Table 1

## Data Records

All raw measurements, the cleaned time series, and the PDF and log files created during the conversion of the raw data to the final database are available in comma-separated format at Zenodo: https://doi.org/10.5281/zenodo.5782922). Online-only table 1 is also stored in the same repository.

## Technical Validation

Visual inspection of the scatter plot of bite force against head width (black dots in Fig. 2b) and all literature data points (orange diamonds) revealed that the literature data lies close to the regression through all data points of our database. This impression is corroborated by the comparison of the allometric slopes of the insect bite force database and the literature data, which yielded no statistically significant difference (ANOVA: *F* = 0.102, *p* = 0.75).

Geographical assessment of the collected animals showed that most species of the insect bite force database were collected in Australia (30.7%), Germany (19.1%), and Panama (16.4%). 23.2% of the species were obtained from breeding cultures. The remaining 10.6% of the species were collected in Greece, Slovenia, France, China, and Denmark. Climate region assessment revealed that most species were collected in temperate (54%) and tropical (43.2%) regions. 2.8% came from dry and continental regions combined (Fig. 3b). We did not consider the original geographic distribution of those species obtained from breeding cultures.

**Fig. 3:**
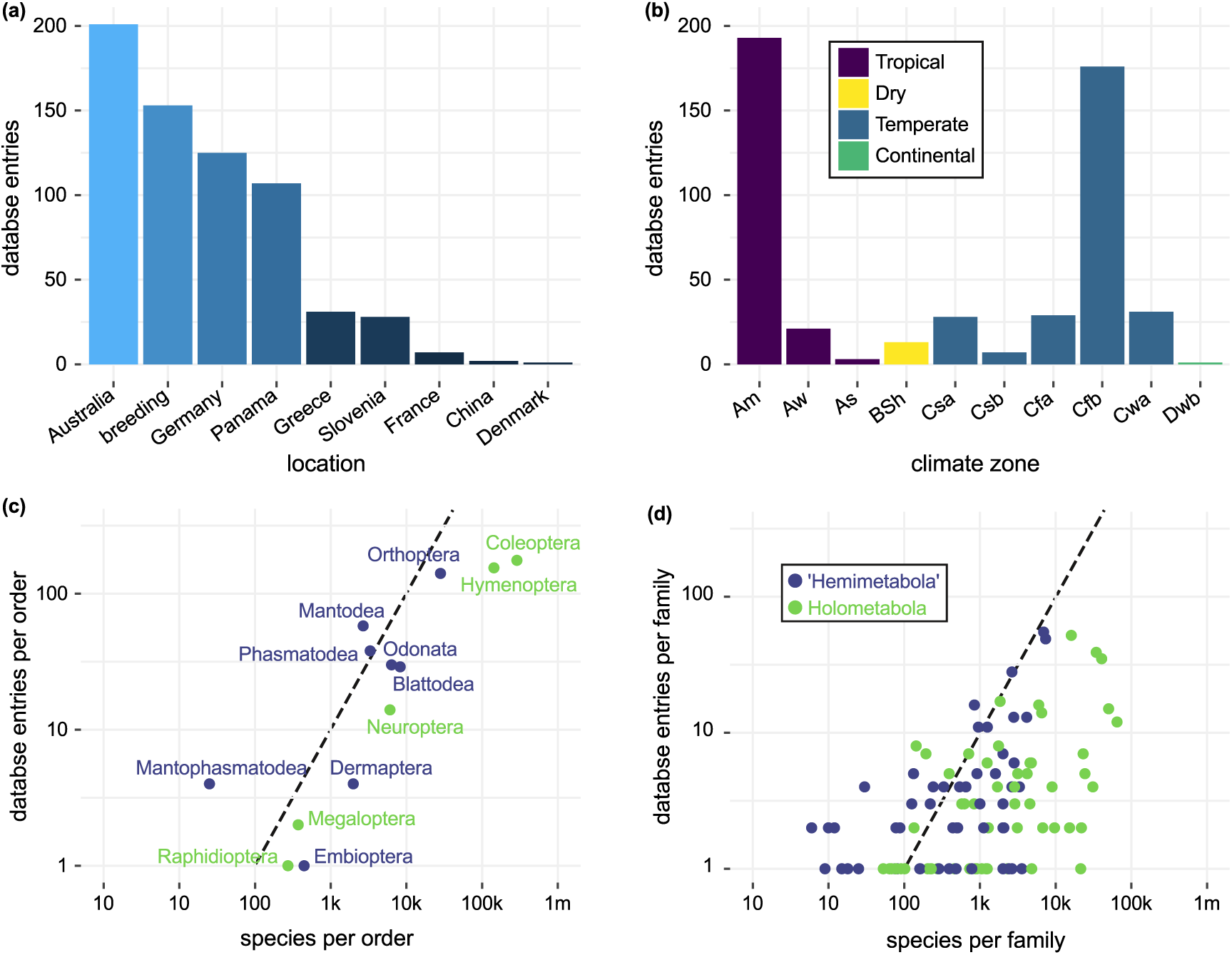
Geographical and phylogenetic coverage of the bite force database. **(a)** Species entries per collection location (country or breeding). **(b)** Species entries per Köppen–Geiger climate zone with specimens sourced from breeding cultures excluded. **(c)** Ratio of database entries compared to species estimated in all insect orders present in the database. **(d)** Ratio of database entries compared to species estimated in all insect families present in the database. The dashed lines in (c,d) mark a ratio of 1 data base entry per 100 estimated species. Abbreviations: **Am**: tropical monsoon; **Aw**: tropical savanna with dry-winter characteristics; **As**: tropical savanna with dry-summer characteristics; **BSh**: semi-arid (steppe) hot; **Csa**: mediterranean hot summer; **Csb**: mediterranean warm/cool summer; **Cfa**: humid subtropical; **Cfb**: oceanic; **Cwa**: dry-winter humid subtropical; **Dwb**: warm summer continental.

A total of 13 biting-chewing insect orders are present in the database (Fig. 3d). We could not obtain life animals from the orders Zoraptera and Grylloblattodea. Bite force measurements of the few species of Plecoptera, Mecoptera, and Trichoptera that were available failed because no voluntary biting could be elicited in these specimens. We did not attempt measuring available representatives of Psocoptera and the biting-chewing “mandibulate archaic moths” (Lepidoptera: Micropterigoidea) due to their minute size. The assessment of phylogenetic coverage of the orders and families showed that most families are represented by less than one species entry per 100 estimated species (Fig. 3c). While orders were sampled in proportion to their taxonomic diversity (Fig. 3d), we were only able to measure at least 1% of the described species in Mantophasmatodea, Phasmatodea and Mantodea (dots left of dashed line in Fig. 3a). Accordingly, bite forces of only a fraction of all insect species were measured so far. Nevertheless, the database exceeds all previous studies combined in species numbers (30-fold in insects, 3.5-fold in amniotes), marking just the beginning of research on this performance trait in the most species-ridge metazoan clade.

## Usage Notes

The forceR package^27^ was used to create the insect bite force database, which contains cleaned measurement time series and maximum bite forces of insects. The same package may be used to expand the scarce knowledge on insect bite forces by tackling questions regarding the evolution of bite lengths, frequencies, and bite curve shapes by semi-automatically extracting individual bite curves from these measurements. Additionally, the maximum bite force values presented in Online-only table 1 can be used for a wide range of in-depth studies on the morphological and ecological predictors and macroevolution of this important performance trait in the megadiverse insects.

## Code Availability

The R code to convert the raw measurements into the final database and to create all tables and figures used in this publication can be found at https://github.com/Peter-T-Ruehr/Insect_Bite_Force_Database.

## Acknowledgements

We would like to express our deep gratitude to the numerous people that helped in collecting (J. Dorey (Flinders University, Adelaide, Australia), E. P. Fagan-Jeffries (University of Adelaide, Adelaide, Australia), L. R. Beische, M. Bläser, R. Predel (University of Cologne, Cologne, Germany), A. M. Jansen, L. Hamann (University of Bonn, Bonn, German), M. Giannotta (Australian National University, Adelaide, Australia), Sam Finnie (University of South Bohemia, Ceske Budejovice, Czech Republic) J. Hamann (Refrath, Germany) and identifying (T. Frenzel (University of Koblenz - Landau, Koblenz, Germany), J. Dorey (Flinders University, Adelaide, Australia), D. C. F. Rentz (James Cook University Cairns Campus, Cairns, Australia), S. You Ning, J. Rodriguez Arrieta (Australian National Insect Collection, Canberra, Australia), E. P. Fagan-Jeffries (University of Adelaide, Adelaide, Australia), M. Giannotta, Y. Living’ Li (Australian National University, Adelaide, Australia), A. Richter (Friedrich-Schiller-Universität Jena, Jena, Germany)) many of the animals used in this study. R. Plarre (Bundesanstalt für Materialforschung und-prüfung (BAM), Berlin, Germany), L. R. Beische, M. Bläser, R. Predel, T. Weihmann (University of Cologne, Cologne, Germany), S. Büsse (Kiel University, Kiel, Germany), W. Hickler (Gelsenkirchen, Germany), C. J. Schwarz (Bochum, Germany), M. Sebesta (Antstore, Berlin, Germany) and B. and D. Drenske (AntsNature, Berlin, Germany) are thanked for providing life animals from their breeding cultures and/or collection trips free of charge. Collections in Australia were carried out at the Daintree Rainforest Observatory (James Cook University, Australia) and under the permit number PTU19-002400 of the Queensland Parks and Wildlife Service. PTR, MF and AB were supported by the European Research Council (ERC) under the European Union’s Horizon 2020 research and innovation program (grant agreement no. 754290, ‘Mech-Evo-Insect’). AB and CE were supported by the Deutsche Forschungsgemeinschaft (DFG) under the Individual Research Grants program (grant agreement no. BL 1355/4-1).

## Author contributions

PTR wrote all code, prepared the figures, lead the dataset creation and drafted the manuscript. PTR and AB conceived the study and refined the manuscript. CE, MF and AB contributed to the dataset creation. All authors reviewed the manuscript and gave final approval for publication.

## Tables

**Online-only Table 1:** Insect bite force database summary. Taxonomic classification, maximum bite force per measurement (iBite), specimen and species (ID), regular and geometric mean bite force per specimen and species, voltage amplification setting, length measurements, collection coordinates, country, and climate zone for each bite fore measurement.

