## Supplementary Figure 1 for "A bite force database of 654 insect species"

### SUPPORTING INFORMATION

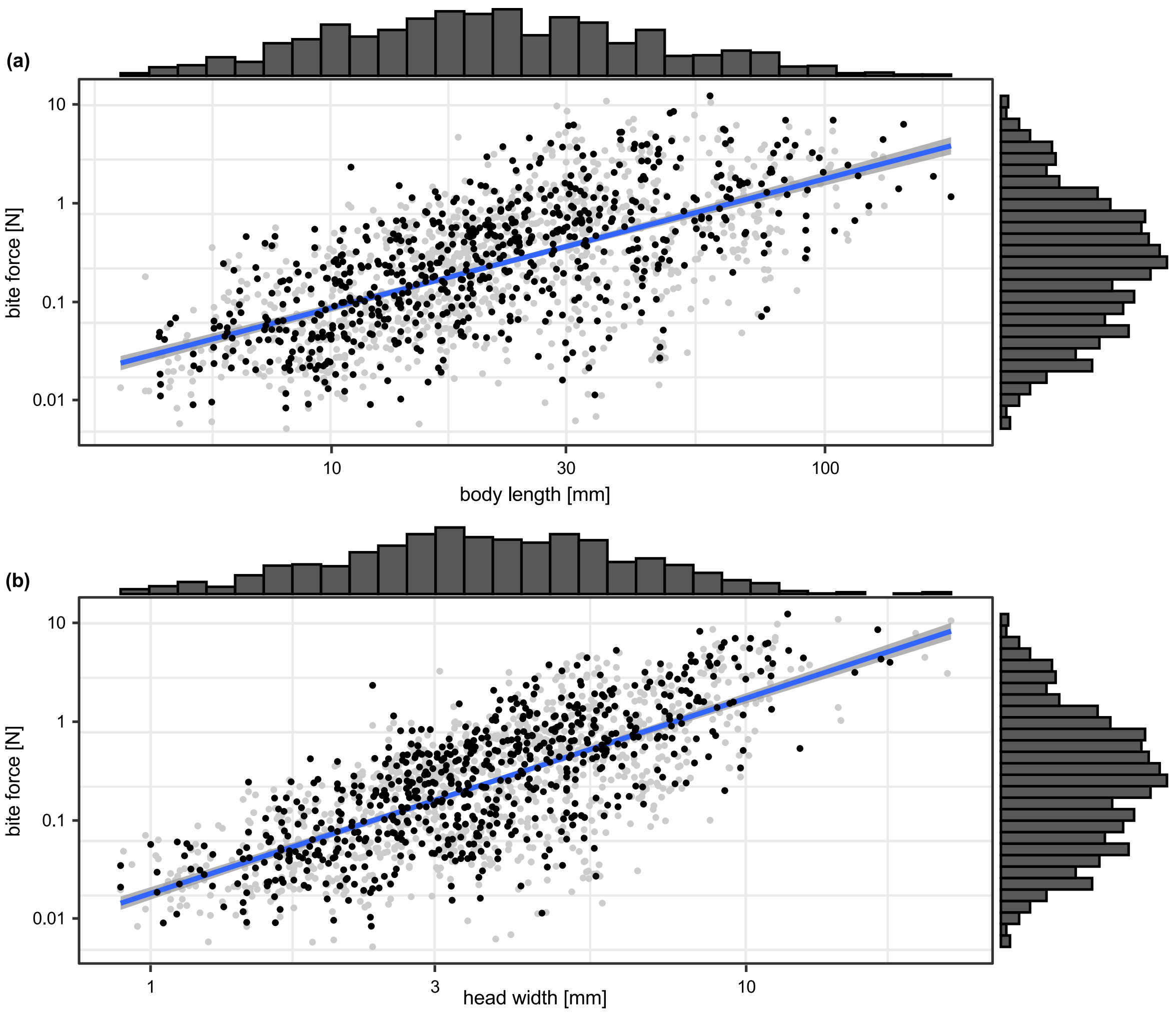


**Supplementary Figure 1:** Maximum bite force against body length (a) and head width (b). Grey dots show means of all maximum bite forces per specimen, black dots show geometric means of all length measurements and maximum bite forces of all specimens per species. Marginal histograms at the x- and y-axes show mean size and mean bite force distribution per specimen, respectively. Regression lines and coefficients refer to log10-linear models of species-wise bite force against body length (a) or head width (b). All axes are log10-transformed.
